# A risk stratification model for lung cancer based on gene coexpression network

**DOI:** 10.1101/179770

**Authors:** Hongyoon Choi, Kwon Joong Na

**Affiliations:** Department of Nuclear Medicine, Cheonan Public Health Center, Chungnam, Republic of Korea; Department of Community Health, Korea Health Promotion Institution, Seoul, Republic of Korea; Department of Clinical Medical Sciences, Seoul National University, College of Medicine, Seoul, Republic of Korea

**Keywords:** lung cancer, gene co-expression network, systems biology, deep learning, microarray

## Abstract

Risk stratification model for lung cancer with gene expression profile is of great interest. Instead of the previously reported models based on individual prognostic genes, we aimed to develop a novel system-level risk stratification model for lung adenocarcinoma based on gene coexpression network. Using multiple microarray datasets obtained from lung adenocarcinoma, gene coexpression network analysis was performed to identify survival-related network modules. Representative genes of these network modules were selected and then, risk stratification model was constructed exploiting deep learning algorithm. The model was validated in two independent test cohorts. Survival analysis using univariate and multivariate Cox regression was performed using the output of the model to evaluate whether the model could predict patients’ overall survival independent of clinicopathological variables. Five network modules were significantly associated with patients’ survival. Considering prognostic significance and representativeness, genes of the two survival-related modules were selected for input data of the risk stratification model. The output of the model was significantly associated with patients’ overall survival in the two independent test sets as well as training set (p < 0.00001, p < 0.0001 and p = 0.02 for training set, test set 1 and 2, respectively). In multivariate analyses, the model was associated with patients’ prognosis independent of other clinical and pathological features. Our study presents a new perspective on incorporating gene coexpression networks into the gene expression signature, and the clinical application of deep learning in genomic data science for prognosis prediction.

## 1. Introduction

Recently, risk stratification based on gene expression profiles is of major biomedical interest in lung cancer research (*1–6*). Previous studies developed risk stratification models that mostly focused on individual prognostic genes. However, these studies have not fully considered the nature of biological networks and their systematic properties. Since it is more evident that biological processes are derived from numerous interactions between many cellular components, gene network analysis could provide valuable information about cancer pathogenesis and therapeutic interventions (*7*). Among the various biological networks, gene co-expression network has some strengths: not relying on prior information about genes, avoiding biologically wrong assumptions about independence of gene expression levels, and alleviating multiple testing problems (*8*).

Lung cancer – mainly, non-small cell lung cancer – is one of the most common cancers and is the leading cause of cancer-related death worldwide (*9, 10*). Currently, TNM staging system is a universal guideline for prognosis prediction and treatment decision. However, heterogeneous molecular features of lung cancer require diverse adjuvant treatment options and lead to different prognosis even in the same stage (*11*). Hence, there has been a constant need for developing better risk stratification models to predict accurate prognosis and to improve cancer-related survival.

The main objectives of this study were 1) to identify survival-related gene co-expression network modules, 2) and to propose a deep learning (DL)-based risk stratification model reflecting survival-related network modules. Using public microarray datasets from the Gene Expression Omnibus (GEO), we identified survival-related network modules of lung adenocarcinoma. Subsequently, we constructed DL-based prognostic score using representative genes of survival-related network modules and it showed great prognostic property in all cohorts.

## 2. Materials and Methods

### 2.1. Gene expression data and preprocessing

Microarray data sets were searched from the National Center for Biotechnology Information GEO database (https://www.ncbi.nlm.nih.gov/geo/) (*12*) using keywords ‘lung cancer’, ‘lung adenocarcinoma’, or ‘adenocarcinoma’. We searched for studies analyzed by single platform (Affymetrix HG-U133A Plus 2.0) in order to obtain high proportion of overlapping genes. In total, eleven microarray data sets were included and raw gene expression data were downloaded from the GEO data repository for preprocessing step (*13–21*). We selected two microarray data sets with survival information (accession number GSE31210 (*20*) and GSE30219 (*21*)) as independent test sets, and the others (*13–19*) with or without survival information as the training set. For those microarray data sets containing multiple histologic types of lung cancer, only the samples from adenocarcinoma were extracted. Detailed information of data source used in this study can be found in **Supplementary Table 1**. All available clinico-pathological variables (age, sex, smoking status, stage, and molecular subtypes) and survival information (survival status and duration) were compiled from each microarray data sets using ‘GEOquery’ package (*22*) (**Table 1)**.

**Table 1.** Demographic and baseline clinical characteristics of patients.

| Variables |  | Training set (n = 533) |  | Test set 1 (GSE31210, n = 226) | Test set 2 (GSE30219, n = 85) |
| --- | --- | --- | --- | --- | --- |
|  |  |  | *Available data* |  |  |
| **Sex** | Female : Male | 229:209 (52.3%:47.7%) | 438 | 121:105 (53.5%:46.5%) | 19:66 (22.4%:77.6%) |
| **Age** |  | 65.46 ± 10.14 | 313 | 59.58 ± 7.40 | 61.49 ± 9.28 |
| **Smoking** | Current | 52 (29.1%) |  | 111(49.1%) |  |
|  | Ex-smoker | 92 (51.4%) | 179 |  |  |
|  | Never | 35 (19.5%) |  | 115 (50.9%) |  |
| **T stage** | T1 | 79 (43.2%) |  |  | 71 (83.5%) |
|  | T2 | 99 (54.1%) |  |  | 12 (14.1%) |
|  | T3 | 4 (2.2%) | 183 |  | 2 (2.4%) |
|  | T4 | 1 (0.5%) |  |  |  |
| **N stage** | N0 | 136 (78.2%) |  |  | 82 (96.5%) |
|  | N1 | 34 (19.5%) | 174 |  | 3 (3.5%) |
|  | N2 | 4 (2.3%) |  |  |  |
| **M stage** | M0 | 127 (100%) | 127 |  | 85 (100%) |
| **Stage** | IA | 91 (29.1%) |  | 114 (50.4%) |  |
|  | IB | 127 (40.6%) |  | 54 (23.9%) |  |
|  | II | 73 (23.3%) | 313 | 58 (25.7%) |  |
|  | III | 18 (5.7%) |  |  |  |
|  | IV | 4 (1.3%) |  |  |  |
| **Mutation** | All negative |  |  | 68 (30.1%) |  |
|  | ALK fusion |  |  | 11 (4.9%) |  |
|  | EGFR mutation |  |  | 127 (56.2%) |  |
|  | KRAS mutation |  |  | 20 (8.8%) |  |
| **Status** | Death : Alive | 106:183 (36.7%:63.3%) | 289 | 35:191 (15.5%:84.5%) | 45:40 (52.9%:47.1%) |
| **Survival time** |  | 52.08 months<br>(0.20 - 190.40) | 289 | 58.150 months<br>(7.37 - 128.80) | 68.00 months<br>(0.00 - 221.00) |

We generated the training set by assembling nine microarray data sets through stepwise preprocessing method described below. The raw gene expression data from microarray data sets were called and normalized using robust multichip average method using the ‘affy’(*23*) package. On a study-by-study basis, we removed invalid and duplicated probe sets by ‘featureFilter’ function in ‘genefilter’ package (*24*) and mapped array probe sets for the respective gene symbols. In addition, to remove poor quality probes, we filtered out probe sets with low expression level (signal intensity < log_2_(100) in at least 5% of samples within at least one study) and low variability (interquartile range < 0.5). As we combined microarray data from different studies, we performed additional normalization using Combat algorithm (*25*) in order to eliminate potential batch effects. Lastly, we detected the outliers by calculating the inter-array correlation based on Pearson’s correlation coefficient for all samples, and removed them. As a result, the training set contained 4615 probe sets from 510 lung adenocarcinoma samples including 273 samples with available survival information.

The raw gene expression data of both test sets were called and normalized as the same method with the training set. One outlier sample was removed from the test set 2; consequently, the test set 1 and 2 included 226 and 84 lung adenocarcinoma samples respectively.

### 2.2. Weighted gene co-expression network construction from the training set

We used weighted gene co-expression network analysis (WGCNA) package (*26, 27*) to build a weighted gene co-expression network from the training set. We created a correlation matrix on the basis of Pearson’s correlation coefficient for all pair-wise genes across all samples. The power –the key parameter for weighted network– was selected to optimize both the scale-free topology and sufficient node connectivity and we chose a threshold of 6 in this study (**Supplementary Fig. 1**). The correlation matrix was transformed into adjacency matrix (matrix of connection strength) using the power function, and pair-wise topological overlap (TO) between genes were calculated. We identified network modules using hierarchical clustering method with TO dissimilarity as the distance measure. The modules were detected using dynamic tree cut algorithm (*28*) in WGCNA package, defining height cutoff value of 0.99, deep split as 2, and minimum module size cutoff value of 30. Genes that were not assigned to any module were classified to color gray (Fig. 1).

**Figure 1.**
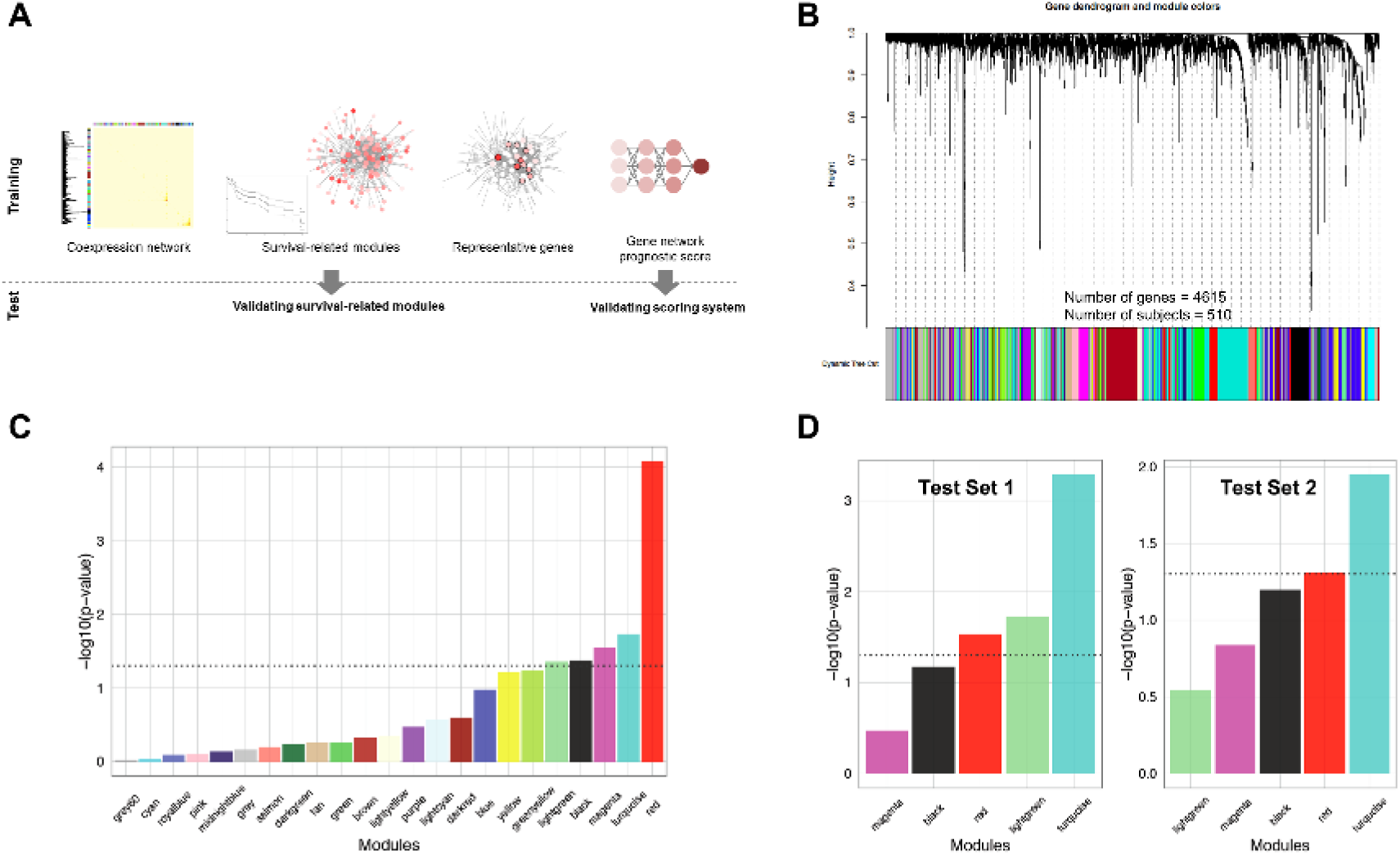
Gene co-expression network construction and survival-related modules identification. (A) A schematic diagram summarizing our risk stratification modeling strategy. Gene co-expression network was constructed from the training set. Gene network modules were extracted based on topological overlap. Survival-related modules were identified from the training set and validated in the two test sets. We selected representative genes from survival-related modules, and built network-based prognostic scoring system using deep learning. (B) Gene dendrogram and modules identified by weighted gene co-expression network analysis from the training set. Modules were labeled with different colors. (C) Univariate Cox regression analysis of module eigengene in the training set was performed. Module eigengene is a representative expression value of genes of each module calculated by the principal component analysis. The dotted line represents cutoff value (p-value = 0.05) for significance, and five modules were identified as survival-related network modules. (D) Survival-related network modules were validated in the two test sets using Cox regression analysis. Three modules from test set 1, and two modules from test set 2 were significantly associated with overall survival.

### 2.3. Identification and validation of survival-related network modules

For each module, we summarized the module expression profile by one representative gene, module eigengene (ME), which is the first principal component of the expression matrix of the corresponding module. We used ME as the representative of each module to evaluate association with overall survival (OS). The survival-related network modules were identified using Cox regression analysis in the training set. For validation, the same genes included in the network construction were extracted from each test set, and assigned to the modules identified in the previous step. ME was calculated based on the expression profile of each test set, and the association between ME and OS was evaluated using Cox regression analysis to see whether the modules identified from the training set are also associated with OS in each test set. The modules with uncorrected p-value under 0.05 were regarded as significant survival-related network modules.

### 2.4. Functional annotation and network visualization of survival-related network modules

The enrichment of the gene ontology terms in each module were evaluated based on the hypergeometric test using ‘clusterProfiler’(*29*) package. The gene ontology biological process terms at false discovery rate under < 0.05 in each survival-related module were regarded as significantly enriched terms. The network of two common survival-related network modules (red and turquoise) was visualized with Cytoscape Software 3.4.0 (*30*).

### 2.5. Representative genes selection for risk stratification model construction

Representative genes of the survival-related network modules were selected to construct risk stratification model. Degree of representativeness of genes in each module was calculated by gene module membership (GMM), a correlation coefficient between gene expression profile and module eigengene. Additionally, the relationship between GMM and prognostic significance (p-value) of an individual gene was tested. Prognostic significance of gene was measured by univariate Cox regression analysis for overall survival. Pearson correlation analysis was performed between GMM and prognostic significance for every gene. We selected top 10 genes according to the GMM from the modules which showed significant correlation between GMM and prognostic significance. Accordingly, expression levels of the selected genes in the same network module were highly correlated to each other, and they could be also highly associated with prognosis because of strong correlation between GMM and prognostic significance. The expression levels of selected genes were used for risk stratification model based on DL.

### 2.6. DL-based risk stratification model

Expression profiles of representative genes were used for the input of the deep learning because they were expected to preserve co-expression patterns and to reflect the systematic properties of survival-related network modules. Convolutional neural network (CNN) was specifically used to extract gene expression patterns of modules. It finally produced gene network prognostic score (NetScore).

DL framework was based on a nonlinear proportional hazard model, which assumed hazard function (λ), a product of a time-dependent baseline hazard function (λ_0_) and a risk function determined by covariates:

*λ*(*x, t*) = *λ*_0_(*t*) × *e*^*h*(*x*)^. Conventional Cox model for the risk stratification using multiple covariates (*x*_1_,*x*_2_, … *x*_*n*_) estimates the risk function *h*(*x*) by a combination of linear functions.

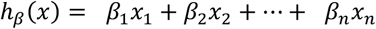

DL-based risk stratification modeling also adopts proportional hazard model, however, replaces linear risk function with the output of neural network (*31*). We designed a simple CNN to estimate risk function, *h*_*θ*_(*x*). Firstly, 1-dimensional convolutional filters were applied. Filter size was same as the input length, 10. Thus, the number of the output of the first layer was same as the number of convolutional filters. Genes in different modules were inputted as different channels. We set the number of filters were 24. The outputs of convolutional layer were hierarchically connected to three fully-connected (FC) layers. Each FC layer had 24 nodes except final output layer. For FC layers, a dropout function was applied to reduce overfitting and learn more robust features. This function randomly drops the connections with predefined probability. We set the probability as 0.5. The final output of CNN, *h*_*θ*_(*x*), was a single node.

The CNN model was trained by the RMSprop algorithm (*32*). The model was optimized to minimize the loss function, negative log partial likelihood.

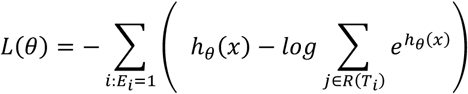

*E*_*i*_=1 represents that the event has occurred in individual *i* at event time *T*_*i*_. *j* ∈ *R*(*T*_*i*_) represents that another patient *j* is still at risk of the event at time *T*_*i*_.

Our framework was trained by initial learning rate with 1x10_-4_ and took 500 epochs for the training. The CNN was implemented using a deep learning library, Keras (ver. 1.0.4) with the Theano (ver. 0.8.2) backend (*33*).

Parameters related to training of the neural network including number of layers, nodes, training epoch and learning rate were determined by 5-fold cross-validation. Training set was randomly divided into 5 subsets. At each step, a single subset was left for testing and other four subsets were used for training. The performance of the model was measured by Harrell’s C-index of the final output score of the model (*34*). The optimal parameters were selected according to the maximum average C-index across the 5-fold of the loop. The predictive value of NetScore was independently validated in two test sets. C-index for each test set was also evaluated.

### 2.7. Comparison of predictability between DL-based model and conventional Cox proportional hazard model

Expression level of all selected genes was fitted into multivariate Cox regression model and the predictive value of the Cox model was evaluated by C-index as in DL-based model. C-index of Cox model was measured by 5-fold cross validation in the training set, and it was calculated in two test sets. C-index of Cox model was compared with that of DL-based model in each cohort (*35*).

### 2.8. Survival analysis using NetScore in all cohorts

Prognostic property of NetScore as continuous variable was evaluated by univariate Cox analysis. To define risk groups, NetScore was dichotomized using the median value in each cohort. Kaplan-Meier method was used to assess survival rates according to the risk groups and survival rate differences were assessed with the log-rank test. Additionally, independent prognostic value of NetScore was assessed by multivariate and subgroup analysis. Multivariate Cox analysis was performed using clinical and pathological variables as well as NetScore. Subgroups were divided on the basis of clinical and pathological features, and univariate Cox analysis of NetScore was performed in each subgroup.

## 3. Results

### 3.1. Gene co-expression network modules from the training set

We aimed at developing a risk stratification model based on gene co-expression networks (Fig. 1A). The networks were constructed from the training set which consists of microarray data of 510 lung adenocarcinoma samples. The clinico-pathological features of all samples from the training set are detailed in Table 1. Using WGCNA, 23 co-expression network modules were identified from the training set (Fig. 1B, **Supplementary Data 1**). The relationship between modules is visualized with hierarchical clustering dendrogram and heatmap of the corresponding ME (**Supplementary Fig. 2**).

### 3.2. Identification of survival-related modules from the training set and validation in test sets

Total five modules were significantly associated with OS (Fig. 1C): red (p < 0.0001), turquoise (p = 0.018), magenta (p = 0.029), black (p = 0.043), and lightgreen (p = 0.044). To validate the survival-related modules, we conducted survival analysis in two independent test sets (GSE31210 as test set 1 and GSE30219 as test set 2; n=226 and 84, respectively). Consequently, turquoise (p = 0.0005), lightgreen (p = 0.019), red (p = 0.030) modules in test set 1, and turquoise (p = 0.011) and red (p = 0.049) modules in test set 2 were significantly associated with OS (Fig. 1D).

The networks of two common survival-related network modules (red and turquoise) are presented in Fig. 2A and 2B. The significantly enriched gene ontology terms of the red module included ‘organic acid catabolic process’, ‘carboxylic acid catabolic process’, ‘small molecule catabolic process’, and the turquoise module included ‘DNA strand elongation involved in DNA replication’, ‘mitotic cell cycle phase transition’, ‘DNA-dependent DNA replication’ (**Supplementary Table 2, Supplementary Data 2**).

**Figure 2.**
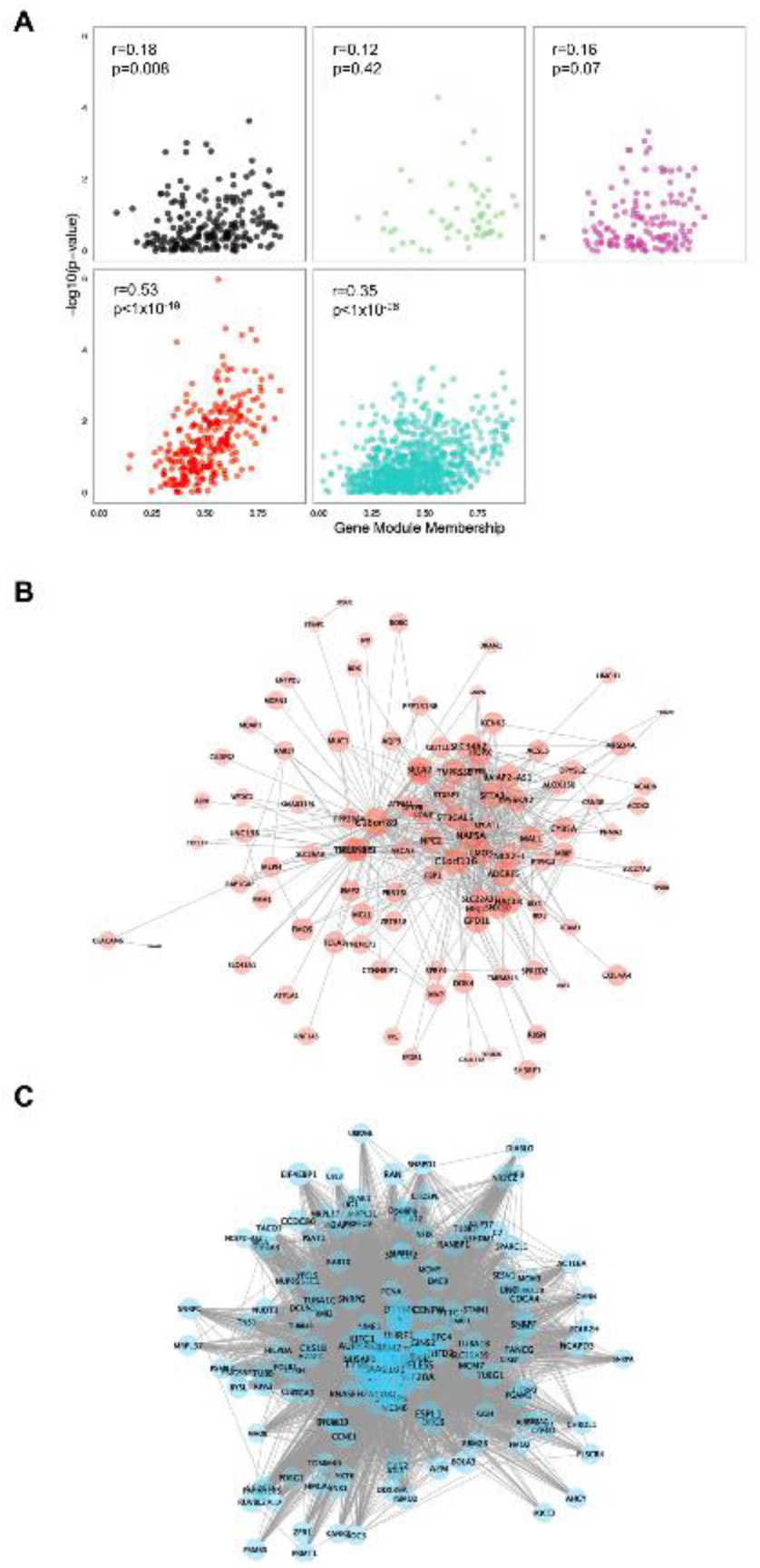
Selection of representative genes of survival-related network modules. Co-expression networks of red (A) and turquoise (B) modules were visualized. Note that 160 genes among 880 genes of turquoise module and their connections were shown. 160 genes were selected according to the gene module membership. Size of nodes is proportional to gene module membership. (C) To construct risk stratification model, representative genes were selected according to the gene module membership. Gene module membership was correlated with the significance of association between individual gene expression and survival. Y-axis represents statistical significance calculated by univariate Cox analysis of individual genes. A strong correlation was found in the red and turquoise modules (r = 0.53 and p < 1x10^-19^ for red module; r = 0.35 and p < 1x10^-23^ for turquoise module).

### 3.3. DL-based risk stratification model using representative genes of survival-related module

By measuring the correlation between gene significance for OS (p-value) and GMM in each survival-related module, we identified two modules demonstrating high correlation with statistical significance (r = 0.53, p < 1x10_-19_ and r = 0.35, p < 1 x 10_-26_ for red and turquoise module, respectively; Fig. 2C). Based on the strong correlation, we could assume that the genes with high representativeness measured by GMM has high significance for OS and are the most important elements of the module; therefore, we selected top 10 genes according to GMM from the red and turquoise modules for the DL-based risk stratification model construction (**Supplementary Fig. 3**).

The expression profiles of selected 20 genes were used as input data of the risk stratification model (Fig. 3A). NetScore, the final output of our model, was significantly associated with OS in the training and two test sets (Fig. 3B) (p < 0.00001, p < 0.0001 and p = 0.02 for training set, test set 1 and 2, respectively). Subjects were divided into two groups, high- and low-risk groups, according to the median value of NetScore in each cohort. The high-risk group was significantly associated with OS in the training set (p < 0.0001; Fig. 3C) and in test set 1 (p < 0.0001; Fig. 3D). A trend of the association was also shown in test set 2 (p = 0.054; Fig. 3E).

**Figure 3.**
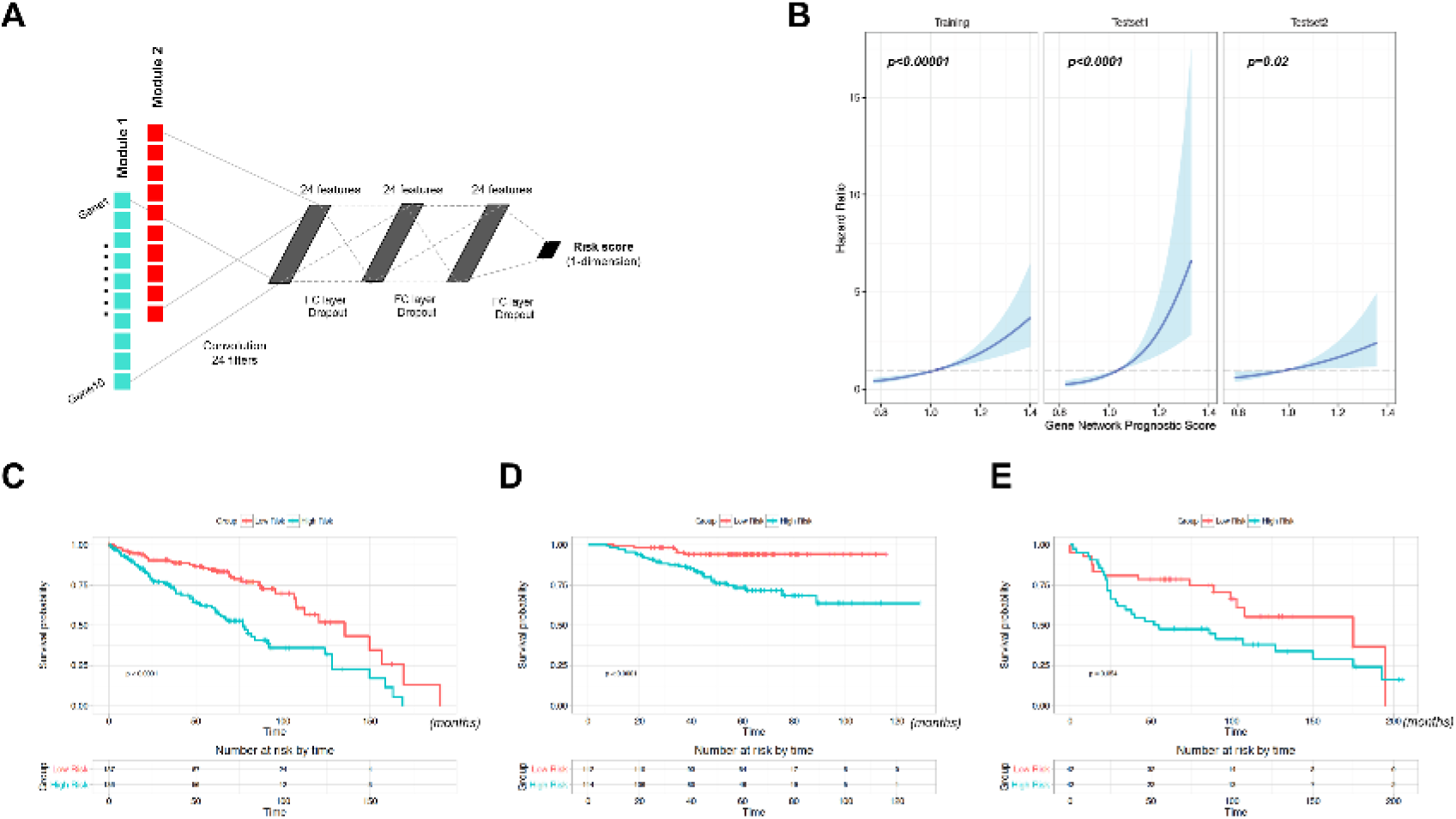
Risk stratification model using representative genes of survival-related network modules. (A) To construct risk stratification model, deep convolutional neural network was used. Input data were expression value of top 10 genes from each of red and turquoise module. The first layer consists of one-dimensional convolutional filters which extract gene expression patterns of each module. Three additional fully-connected (FC) layers were followed and connected to the output score gene network prognostic score (NetScore). (B) Univariate Cox regression analysis of NetScore as a continuous variable was performed in the training and two test sets. It shows significant association between the score and overall survival in all sets. The blue line represents hazard ratio for overall survival and the blue area represents 95% confidence interval. (C-E) Overall survival of dichotomized group according to NetScore was depicted by Kaplan-Meier survival curve. The statistical difference was tested by log-rank test. The high risk group showed worse survival in the training set (C) and test set 1 (D) with statistical significance. The high risk group of the test set 2 (E) also showed worse prognosis though the difference did not reach statistical significance.

### 3.4. NetScore as an independent predictive factor for prognosis

Cox multivariate analysis revealed that the risk group was associated with OS independent of stage as well as other clinico-pathological features in the training set and test set 1 (Table 2). The independent predictive factors for OS in Cox multivariate analysis were the risk group (p = 0.001) and T-stage 3 (p = 0.030) in training set, and the risk group (p = 0.01) and EGFR mutation status (p = 0.005) in test set 1. In the test set 2, there was no feature significantly associated with OS in univariate Cox analysis, though the high risk group showed a trend of unfavorable prognosis (p = 0.06). We also evaluated the prognostic value of NetScore in subgroups divided by clinical and pathological features. In the training set, the high risk group was significantly associated with poor prognosis in subgroups regardless of age and T-stage. In all subgroups, a trend of close relationship between the risk group and OS was found except never-smoking subgroup (Fig. 4A, **Supplementary Fig. 4**). According to subgroup analysis in test set 1, the risk group was closely associated with OS in male, old-aged, ever/never smokers, stage IA/IB, EGFR positive and all negative mutation subgroups (Fig. 4B, **Supplementary Fig. 5**). A trend of association between the risk group and OS was also revealed in each subgroup of test set 2, regardless of clinical features including sex, age and T stage (Fig. 4C, **Supplementary Fig. 6**).

**Table 2.** Univariate and multivariate Cox regression analysis of the risk stratification model and clinicopathological variables for overall survival in the training and test sets.

| Variables | Univariate analysis |  | Multivariate analysis |  |
| --- | --- | --- | --- | --- |
|  | Hazard Ratio (95% CI) | p-value | Hazard Ratio (95% CI) | p-value |
| **Training set** |  |  |  |  |
| Gene network prognostic score - High risk group | 2.51 (1.65 - 3.81) | < 0.001 | 3.057 (1.556 - 6.006) | 0.001 |
| Age - Older than 60 | 1.66 (0.96 - 2.86) | 0.068 |  |  |
| Sex – Male | 1.28 (0.86 - 1.88) | 0.221 |  |  |
| Smoking status - Ex-smoker | 0.59 (0.31 - 1.12) | 0.108 |  |  |
| Smoking status - Never-smoker | 0.51 (0.22 - 1.19) | 0.120 |  |  |
| T stage : II | 2.50 (1.31 - 4.79) | 0.006 | 1.266 (0.599 - 2.674) | 0.537 |
| T stage : III | 13.32 (2.89 - 61.32) | 0.001 | 5.895 (1.189 - 29.237) | 0.030 |
| N stage : I | 2.27 (1.28 - 4.05) | 0.005 | 1.762 (0.943 - 3.294) | 0.076 |
| **Test set 1** |  |  |  |  |
| Gene network prognostic score - High risk group | 4.39 (1.92-10.06) | 0.0004 | 2.97 (1.25-7.09) | 0.01 |
| Age - Older than 60 | 1.27 (0.65-2.48) | 0.49 |  |  |
| Sex – Male | 1.52 (0.78 - 2.96) | 0.22 |  |  |
| Smoking status - Never-smoker | 0.61 (0.31-1.19) | 0.15 |  |  |
| Stage: II | 4.23 (2.17 - 8.24) | 0.00002 |  |  |
| EGFR mutation + | 0.47 (0.24-0.93) | 0.03 | 2.74 (1.36-5.54) | 0.005 |
| KRAS mutation + | 0.87 (0.27-2.85) | 0.82 | 0.64 (0.32-1.27) | 0.20 |
| **Test set 2** |  |  |  |  |
| Gene network prognostic score - High risk group | 1.81(0.98-3.36) | 0.06 |  |  |
| Age - Older than 60 | 1.33(0.73-2.43) | 0.35 |  |  |
| Sex – Male | 0.83(0.40-1.75) | 0.63 |  |  |
| Stage: T2 | 1.65(0.84-3.25) | 0.14 |  |  |

**Figure 4.**
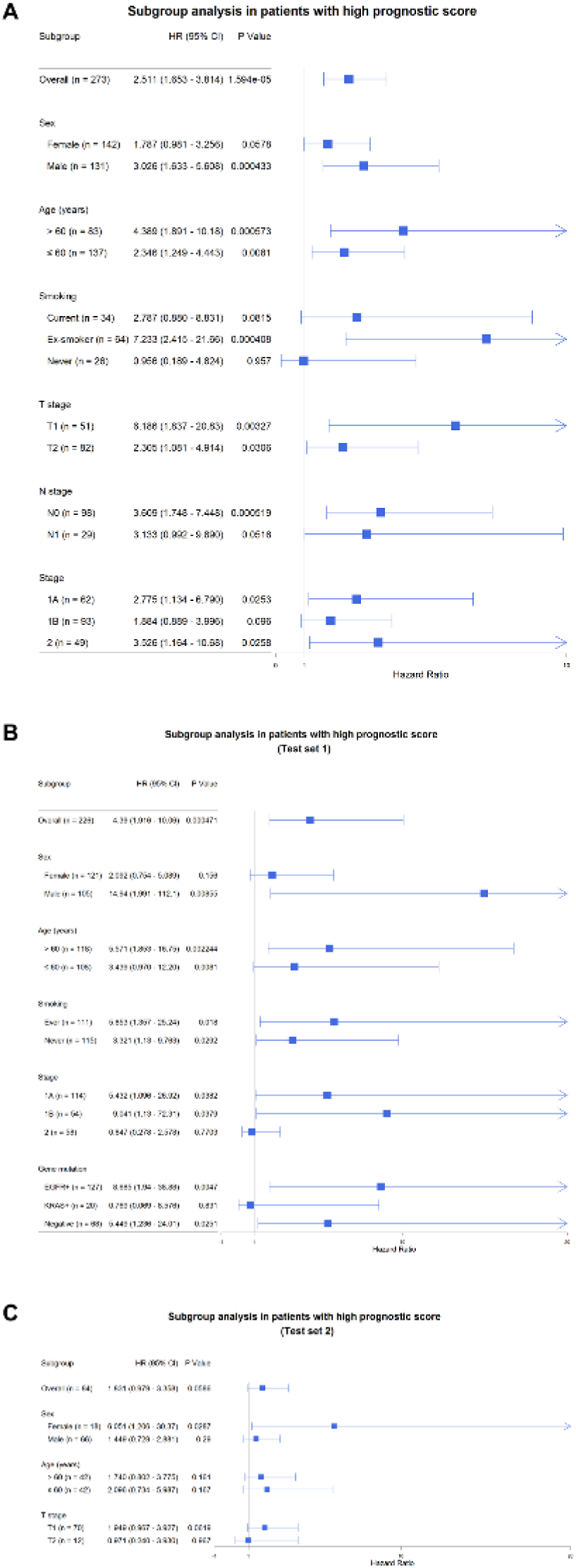
Subgroup analysis using NetScore. (A) Predictive value of our risk stratification model was tested in subgroups classified by clinico-pathological characteristics of the training set. A trend of association between the risk group and overall survival was found in all subgroups. (B, C) The same subgroup survival analysis was also performed in both test sets. (B) The risk group was associated with overall survival regardless of clinico-pathological variables except female, stage II and KRAS mutation subgroups in test set 1. (C) Regardless of subgroups, a trend of poor prognosis in high risk group was also found in the test set 2

## 4. Discussion

In this study, we developed a risk stratification model for lung adenocarcinoma based on gene co-expression networks and deep learning. Survival-related network modules were identified in multiple cohorts and representative genes of these modules were selected for risk stratification modeling. The model constructed by deep CNN reflects gene expression patterns of survival-related network modules and it provides prognostic score, NetScore. The NetScore was significantly associated with OS in all cohorts and also an independent predictor for OS from clinico-pathological variables.

The model based on survival-related network modules can provide more robust risk stratification compared with models focusing on statistical combination of individual prognostic genes which have been proposed in the previous studies (*1–6*). In spite of their promising results, several models failed to validate in independent samples of other study (*4*). Furthermore, there were few overlapping significant prognostic genes in the previous models. A meta-analysis of published gene expression data revealed that few genes were associated with survival of lung adenocarcinoma (*36*). The result of few significant prognostic genes in large samples implied the limitation of usage of individual genes for risk stratification. Besides, selection of individual significant genes has a substantial problem of multiple statistical testing (*37*). Instead of these previous approaches, systemic approach integrating gene interaction as well as individual genes would be a breakthrough for robust risk stratification modeling because variation patterns of their expression levels can be associated with prognosis.

Functional annotation elucidated the role of specific gene network in lung adenocarcinoma pathophysiology. The red module was functionally associated with catabolic process of organic acid and carboxylic acid. As cancer cells rely on aerobic glycolysis and facilitate the metabolic process of amino acid, nucleotides and lipid for rapid proliferation, genes related to fatty-acid and amino acid metabolism could reflect progression of cancer cell (*38*). The turquoise module was related to DNA replication and cell-cycle. A previous gene co-expression analysis also revealed that cell-cycle related genes were closely associated with lung cancer prognosis (*39*). Furthermore, overexpression of cyclins has repeatedly been associated with poor prognosis in lung cancer (*40*). Our result also emphasizes the importance of cell-cycle genes in lung cancer prognosis.

Recently, DL has dramatically improved data analysis in genomics and imaging fields (*41, 42*). The main contribution of DL for our risk stratification model is firstly applying convolutional neural network to gene expression data. It was used for extracting multiple gene expression patterns by applying convolutional filters. Another contribution is to solve regression problems of survival data by using a specialized loss function (*31*). We compared predictive accuracy of DL-based model and conventional Cox proportional hazard model obtained from the expression level of selected 20 genes. Predictability of the DL-based model was significantly higher than that of the Cox model in test set 1 (C-index= 0.709±0.042 and 0.608±0.046, respectively; p = 0.004). It was also higher in the training set and test set 2 though the difference did not reach statistical significance (**Supplementary Fig. 7**). To our knowledge, NetScore is the first study that apply deep convolutional neural network to high-dimensional gene expression data for predicting prognosis. By applying this novel approach to various genomic data, risk stratification and survival prediction could be improved compared with conventional Cox model.

NetScore was trained by various samples with different clinico-pathological characteristics. We found NetScore was associated with sex, smoking status, stage and molecular subtypes (**Supplementary Fig. 8**). Briefly, a trend of high NetScore was found in male, smokers, late stage and KRAS mutation positive samples. Nonetheless, NetScore was significantly associated with OS independent of clinico-pathological variables according to multivariate and subgroup analyses. Of note, NetScore was significant predictor in early stage subgroups (stage IA/IB). This finding could be important because the new risk stratification could identify patients who might need adjuvant chemotherapy. For example, a recent clinical trial using 15-gene signature based on individual prognostic genes showed successful selection of patients with stage IB and II NSCLC who would most likely benefit from adjuvant chemotherapy (*43*). In the future, as a new prognostic biomarker based on gene network, the usefulness of NetScore should be tested whether it could affect clinical decision, and compared with the previous prognostic models using individual genes.

## 5. Conclusion

We developed a risk stratification model for lung adenocarcinoma using gene co-expression network. A future extension of our work would be to apply this approach to the co-expression networks of other cancer types. In terms of technical improvement, modification of DL architecture and selection process of representative genes could improve the prediction accuracy. Finally, we expected that a prospectively designed clinical trial with well-controlled clinico-pathological variables would help find clinical application of our new risk stratification model.

## Acknowledgements

This research did not receive any specific grant from funding agencies in the public, commercial, or not-for-profit sectors.

## REFERENCES

1. H. Y. Chen et al., A five-gene signature and clinical outcome in non-small-cell lung cancer. The New England journal of medicine 356, 11–20 (2007).

2. Y. Xie et al., Robust gene expression signature from formalin-fixed paraffin-embedded samples predicts prognosis of non-small-cell lung cancer patients. Clinical cancer research: an official journal of the American Association for Cancer Research 17, 5705–5714 (2011).

3. M. Skrzypski et al., Three-gene expression signature predicts survival in early-stage squamous cell carcinoma of the lung. Clinical cancer research: an official journal of the American Association for Cancer Research 14, 4794–4799 (2008).

4. A. Director's Challenge Consortium for the Molecular Classification of Lung et al., Gene expression-based survival prediction in lung adenocarcinoma: a multi-site, blinded validation study. Nature medicine 14, 822–827 (2008).

5. P. Roepman et al., An immune response enriched 72-gene prognostic profile for early-stage non-small-cell lung cancer. Clinical cancer research: an official journal of the American Association for Cancer Research 15, 284–290 (2009).

6. P. C. Boutros et al., Prognostic gene signatures for non-small-cell lung cancer. Proceedings of the National Academy of Sciences of the United States of America 106, 2824–2828 (2009).

7. A. L. Barabasi, N. Gulbahce, J. Loscalzo, Network medicine: a network-based approach to human disease. Nature reviews. Genetics 12, 56–68 (2011).

8. B. Zhang, S. Horvath, A general framework for weighted gene co-expression network analysis. Stat Appl Genet Mol Biol 4, Article17 (2005).

9. R. L. Siegel, K. D. Miller, A. Jemal, Cancer statistics, 2015. CA: a cancer journal for clinicians 65, 5–29 (2015).

10. L. A. Torre et al., Global cancer statistics, 2012. CA: a cancer journal for clinicians 65, 87–108 (2015).

11. W. Pao, N. Girard, New driver mutations in non-small-cell lung cancer. The lancet oncology 12, 175–180 (2011).

12. T. Barrett et al., NCBI GEO: archive for functional genomics data sets—10 years on. Nucleic acids research 39, D1005–D1010 (2011).

13. S. D. Der et al., Validation of a histology-independent prognostic gene signature for early-stage, non–small-cell lung cancer including stage IA patients. Journal of Thoracic Oncology 9, 59–64 (2014).

14. J. Hou et al., Gene expression-based classification of non-small cell lung carcinomas and survival prediction. PloS one 5, e10312 (2010).

15. J. Botling et al., Biomarker discovery in Non–Small cell lung cancer: Integrating gene expression profiling, meta-analysis, and tissue microarray validation. Clinical Cancer Research 19, 194–204 (2013).

16. R. Kuner et al., Global gene expression analysis reveals specific patterns of cell junctions in non-small cell lung cancer subtypes. Lung cancer 63, 32–38 (2009).

17. P. Micke et al., Gene copy number aberrations are associated with survival in histologic subgroups of non-small cell lung cancer. Journal of thoracic oncology 6, 1833–1840 (2011).

18. A. C. Borczuk et al., Progression of Human Bronchioloalveolar Carcinoma to Invasive Adenocarcinoma Is Modeled in a Transgenic Mouse Model of K-ras–Induced Lung Cancer by Loss of the TGF-β Type II Receptor. Cancer research 71, 6665–6675 (2011).

19. L. Ding et al., Somatic mutations affect key pathways in lung adenocarcinoma. Nature 455, 1069–1075 (2008).

20. H. Okayama et al., Identification of genes upregulated in ALK-positive and EGFR/KRAS/ALK-negative lung adenocarcinomas. Cancer research 72, 100–111 (2012).

21. S. Rousseaux et al., Ectopic activation of germline and placental genes identifies aggressive metastasis-prone lung cancers. Science translational medicine 5, 186ra166–186ra166 (2013).

22. S. Davis, P. S. Meltzer, GEOquery: a bridge between the Gene Expression Omnibus (GEO) and BioConductor. Bioinformatics 23, 1846–1847 (2007).

23. L. Gautier, L. Cope, B. M. Bolstad, R. A. Irizarry, affy—analysis of Affymetrix GeneChip data at the probe level. Bioinformatics 20, 307–315 (2004).

24. R. Gentleman, V. Carey, W. Huber, F. Hahne, Genefilter: methods for filtering genes from high-throughput experiments. R package version 1, (2015).

25. W. E. Johnson, C. Li, A. Rabinovic, Adjusting batch effects in microarray expression data using empirical Bayes methods. Biostatistics 8, 118–127 (2007).

26. P. Langfelder, S. Horvath, WGCNA: an R package for weighted correlation network analysis. BMC bioinformatics 9, 1 (2008).

27. B. Zhang, S. Horvath, A general framework for weighted gene co-expression network analysis. Statistical applications in genetics and molecular biology 4, 1128 (2005).

28. P. Langfelder, B. Zhang, S. Horvath, Defining clusters from a hierarchical cluster tree: the Dynamic Tree Cut package for R. Bioinformatics 24, 719–720 (2008).

29. G. Yu, L.-G. Wang, Y. Han, Q.-Y. He, clusterProfiler: an R package for comparing biological themes among gene clusters. Omics: a journal of integrative biology 16, 284–287 (2012).

30. M. S. Cline et al., Integration of biological networks and gene expression data using Cytoscape. Nature protocols 2, 2366–2382 (2007).

31. J. Katzman et al., Deep Survival: A Deep Cox Proportional Hazards Network. arXiv preprint arXiv:1606.00931, (2016).

32. T. Tieleman, G. Hinton, Lecture 6.5-rmsprop: Divide the gradient by a running average of its recent magnitude. COURSERA: Neural Networks for Machine Learning 4, (2012).

33. F. Bastien et al., Theano: new features and speed improvements. arXiv preprint arXiv:1211.5590, (2012).

34. F. E. Harrell, K. L. Lee, R. M. Califf, D. B. Pryor, R. A. Rosati, Regression modelling strategies for improved prognostic prediction. Statistics in medicine 3, 143–152 (1984).

35. L. Kang, W. Chen, N. A. Petrick, B. D. Gallas, Comparing two correlated C indices with right-censored survival outcome: a one-shot nonparametric approach. Stat Med 34, 685–703 (2015).

36. B. Gyorffy, P. Surowiak, J. Budczies, A. Lanczky, Online survival analysis software to assess the prognostic value of biomarkers using transcriptomic data in non-small-cell lung cancer. PLoS One 8, e82241 (2013).

37. D. B. Allison, X. Cui, G. P. Page, M. Sabripour, Microarray data analysis: from disarray to consolidation and consensus. Nature reviews. Genetics 7, 55–65 (2006).

38. M. G. Vander Heiden, L. C. Cantley, C. B. Thompson, Understanding the Warburg effect: the metabolic requirements of cell proliferation. Science 324, 1029–1033 (2009).

39. Y. Li et al., Network-based approach identified cell cycle genes as predictor of overall survival in lung adenocarcinoma patients. Lung Cancer 80, 91–98 (2013).

40. S. Singhal, A. Vachani, D. Antin-Ozerkis, L. R. Kaiser, S. M. Albelda, Prognostic implications of cell cycle, apoptosis, and angiogenesis biomarkers in non-small cell lung cancer: a review. Clinical cancer research: an official journal of the American Association for Cancer Research 11, 3974–3986 (2005).

41. C. Angermueller, T. Parnamaa, L. Parts, O. Stegle, Deep learning for computational biology. Molecular systems biology 12, 878 (2016).

42. Y. LeCun, Y. Bengio, G. Hinton, Deep learning. Nature 521, 436–444 (2015).

43. C. Q. Zhu et al., Prognostic and predictive gene signature for adjuvant chemotherapy in resected non-small-cell lung cancer. Journal of clinical oncology: official journal of the American Society of Clinical Oncology 28, 4417–4424 (2010).

